# An Integrated Single-Cell and Epigenomic Resource for Comparative Analysis of the Basal Ganglia

**DOI:** 10.64898/2026.01.29.702575

**Authors:** Wenjin Zhang, Wubin Ding, Kai Li, Lei Chang, Amit Klein, Cindy Tatiana Báez-Becerra, Jonathan A. Rink, Anna Bartlett, Huaming Chen, Natalie Schenker, Nelson Johansen, Tyler Mollenkopf, Yuanyuan Fu, Xie Yang, Shane Liu, Chanrung Seng, Benpeng Miao, Tianjie Liu, Quan Zhu, Rebecca D. Hodge, Trygve E. Bakken, Ed S. Lein, Michael Hawrylycz, Xiangmin Xu, M. Margarita Behrens, Bing Ren, Joseph R. Ecker, Ting Wang, Daofeng Li

## Abstract

The basal ganglia regulate motor, cognitive, and affective behaviors, and their dysfunction underlies diverse neurological and psychiatric disorders. Comprehensive, accessible multi-omics resources are needed to understand the regulatory mechanisms governing basal ganglia cell types. Here we present an open, interactive web-based platform for exploring single-cell multi-omics datasets from basal ganglia, generated using 10X Multiome, snm3C-seq, and Paired-Tag technologies from the BICAN (NIH BRAIN Initiative Cell Atlas Network) consortium. The platform is available at https://basalganglia.epigenomes.net/ and enables integrated visualization of gene expression, chromatin accessibility, DNA methylation, histone modifications, and chromatin conformation across cell types and human, macaque, marmoset, and mouse species, with direct genome browser support and comparative epigenomic functionality. Representative analyses demonstrate cell-type-specific regulatory landscapes, conserved and species-specific regulatory elements, and links between epigenomic regulation and transcription. This resource provides a scalable, community-oriented foundation for advancing basal ganglia biology and interpreting regulatory mechanisms relevant to brain function and disease.

**Highlights:**

- Integrated single-cell epigenomic resource for basal ganglia
- Interactive genome browser enables multi-omics and cross-species exploration
- Reveals cell-type-specific and species-specific regulatory landscapes
- Supports community access to complex brain epigenomic datasets

## Introduction

To address the complexity of the human brain, the NIH BRAIN Initiative has established coordinated, large-scale efforts to map brain cell types and their molecular features systematically. The Brain Initiative Cell Atlas Network (BICAN) is a central component of this strategy, aiming to generate a comprehensive, multimodal reference atlas of human and model-organism brain cell types. BICAN integrates transcriptomic, epigenomic, anatomical, and functional measurements across multiple brain regions and species, emphasizing standardized data generation, open data sharing, and cross-consortium interoperability. These principles are intended to enable broad reuse of data resources and to support community-driven discovery^1^.

The basal ganglia (BG) serve as a central hub for integrating inputs from the cortex, thalamus, and brainstem, coordinating these signals to guide adaptive motor, cognitive, and affective behaviors. Disruptions within these circuits are associated with a wide range of neurological and psychiatric conditions^2-4^. Beyond their translational relevance, the basal ganglia were prioritized by the BICAN consortium due to their conserved structure and functional organization across species, making them an ideal system for cross-species comparative studies and for linking molecular and cellular findings to evolutionary and functional principles.

Advances in single-cell and single-nucleus technologies have been instrumental in enabling these efforts^5^. Single-cell RNA sequencing (scRNA-seq)^6^ has revealed extensive transcriptional diversity within neuronal and non-neuronal populations, redefining cell-type classification in the brain^7^. In parallel, single-cell ATAC-seq has enabled the systematic mapping of chromatin accessibility landscapes, providing insights into cis-regulatory elements and transcription factor networks that underlie cell identity and state^8^. Together, these approaches have become foundational tools for constructing cell atlases and for linking molecular profiles to brain function and disease.

More recently, multimodal epigenomic technologies have expanded the scope of single-cell profiling. Single-nucleus methylome and chromatin conformation sequencing (snm3C-seq) enables joint measurement of DNA methylation and three-dimensional genome organization within individual nuclei, providing an integrated view of epigenetic regulation and higher-order chromatin architecture^9^. Paired-Tag technology enables simultaneous profiling of histone modifications and gene expression within the same cells, directly linking chromatin state to transcriptional output^10,11^. Such multimodal measurements are essential for understanding how epigenetic mechanisms shape cell-type-specific gene regulation and how these processes vary across brain regions and species.

Epigenetic regulation plays a critical role in basal ganglia development, neuronal subtype specification, and functional specialization. Cell-type- and region-specific patterns of chromatin accessibility, DNA methylation, histone modifications, and chromatin interactions contribute to regulatory programs that are not fully captured by transcriptomic data alone. Integrating these layers is therefore essential for constructing a comprehensive molecular framework of basal ganglia biology and for interpreting disease-associated genetic variation^12-14^.

While large-scale efforts such as BICAN have generated extensive multi-omics datasets, effective data exploration and integration remain significant challenges for the broader research community. Web-based platforms that provide standardized access, intuitive navigation, and interactive visualization are critical for maximizing the utility of these resources. By enabling users to explore cell types, genomic features, and regulatory modalities within a unified framework and by supporting comparative analyses across species, such portals facilitate hypothesis generation, cross-study integration, and collaborative research. These capabilities are particularly valuable for advancing a systems-level understanding of basal ganglia function and dysfunction.

## Results

### Integrated multi-omics basal ganglia resource and data overview

To enable systematic exploration of basal ganglia molecular architecture, we assembled and integrated a comprehensive collection of single-cell and single-nucleus multi-omics datasets spanning major basal ganglia subregions based on the BICAN cross-species consensus basal ganglia taxonomy^15^. The resource includes transcriptomic, chromatin accessibility, DNA methylation, histone modification, and chromatin conformation data generated using 10X Multiome (scRNA-seq and scATAC-seq^15^), snm3C-seq (Ding et al., co-submit), and Paired-Tag (Chang et al., co-submit) technologies. These datasets capture both neuronal and non-neuronal cell populations and provide complementary views of gene regulation across multiple epigenetic layers.

All datasets were uniformly processed using standardized pipelines to ensure consistency across assays, regions, and species. Cell-type annotations were harmonized across modalities using established marker genes and cross-modality integration strategies, enabling coherent comparisons between transcriptomic and epigenomic profiles. Data are organized by consensus cell taxonomy either on subclass (Supplemental Table 1) or group level (Supplemental Table 2), assay modality, and species, supporting both targeted queries and broad exploratory analyses.

To facilitate accessibility and reuse, the integrated datasets are hosted within a web-based portal that provides structured navigation across cell types and assays. Users can interactively explore data through dynamically generated tables and genome browser views, enabling rapid inspection of regulatory features, including chromatin accessibility, DNA methylation, histone modifications, and chromatin interactions, within defined cell populations. The portal provides a genome browser anchored workflow that guides users from cell-type selection to integrated multi-omics visualization (Figure 1). Users first define cell populations using a hierarchical taxonomy, then select assay modalities of interest, after which the corresponding datasets are dynamically rendered in an interactive genome browser environment. This workflow enables coordinated inspection of chromatin accessibility, histone modifications, DNA methylation, transcriptional activity, and regulatory annotations within a unified genomic context, facilitating both exploratory and hypothesis-driven analyses.

**Figure 1.**
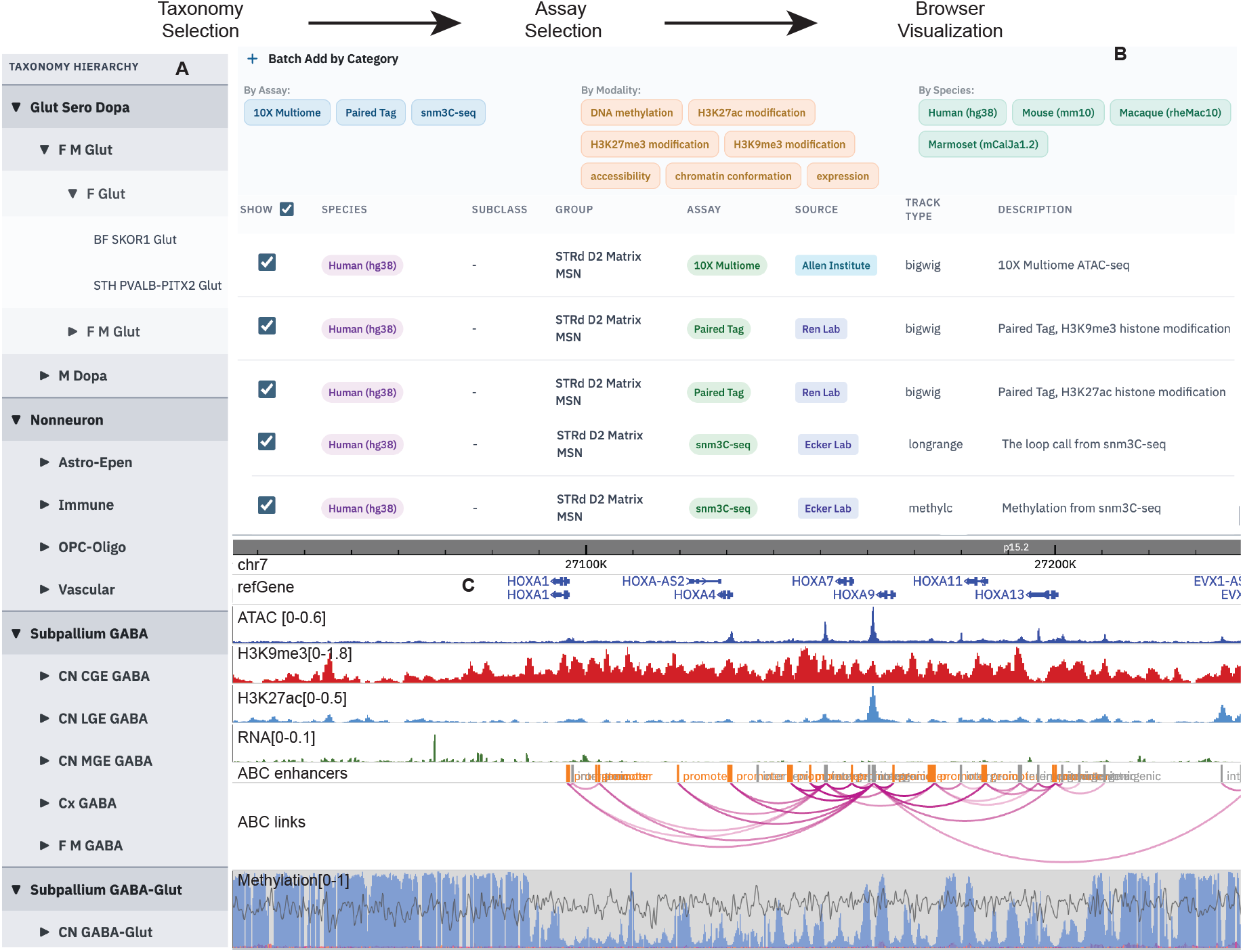
Genome browser anchored visualization workflow of the portal. (A) Data exploration begins with cell-type selection using a hierarchical taxonomy presented in the first tab of the home page, allowing selection at both subclass and group levels. (B) In the second tab, users select assay modalities for visualization. All available assays are preselected by default, with search and filtering tools provided to customize the selection. (C) Based on the selected cell types and assays, the portal renders an interactive genome browser view displaying the corresponding datasets. Standard genome browser operations, including zooming, panning, and gene- or coordinate-based searches, support detailed examination of genomic and epigenomic features. ATAC-seq, histone modification, and RNA-seq tracks are displayed using CPM (counts per million) normalization, consistent across all figures. Activity-by-contact (ABC) enhancers and enhancer-promoter links are derived from Ding et al. (co-submit). DNA methylation data are shown in methylC format^16^, with blue bars representing CpG methylation levels (range 0-1) and gray lines indicating read density.

This integrated data framework establishes a unified entry point for basal ganglia multiomics data and provides the foundation for downstream analyses of cell-type-specific regulatory mechanisms and disease-relevant genomic features.

### Web portal architecture and interactive visualization framework

To support efficient exploration of complex, multimodal datasets, the web-based portal was developed with a modular architecture optimized for scalability, interoperability, and user-driven analysis. The architecture is designed to accommodate continued data expansion as additional assays become available through ongoing consortium and community efforts. The complete list of available data is stored in a TSV (Tab-Separated Values) file that can be updated for future expansion (Supplementary File 1).

The user interface emphasizes intuitive navigation and flexible data access. Users can browse datasets by cell taxonomy, assay modality, or species. One central feature of the portal is its integration with the WashU Epigenome Browser^17,18^, which enables high-resolution, genome-based visualization of multiple epigenomic and transcriptomic tracks. Users can generate customized visualization panels that combine data across assays, cell types, and species. Visualization settings, including track selection, scaling, and synchronization state, can be saved and shared with collaborators, supporting reproducibility and collaborative workflows.

In addition, the resource incorporates cross-species datasets generated within the BICAN consortium, enabling comparative analyses between human and non-human primate (NHP) basal ganglia cell types. It supports cross-species comparative epigenomic analyses by enabling synchronized visualization of homologous genomic regions across multiple species. Using the *FOXP2* locus as an illustrative example, users can simultaneously examine chromatin accessibility and regulatory signatures from human, mouse, macaque, and marmoset basal ganglia datasets (Figure 2). The *FOXP2* gene (forkhead box P2), located on chromosome 7, is a critical transcription factor regulating brain development, specifically in areas managing motor control, such as the basal ganglia and cerebellum^19^. Alignment of epigenomic tracks across species is achieved using genomeAlign tracks^20^ derived from pairwise whole-genome sequence alignments, enabling direct comparison of regulatory features despite differences in genome organization. At the *FOXP2* locus, this approach reveals conserved chromatin accessible regions that are shared across all four species, consistent with evolutionarily conserved enhancer elements. The ability to identify such conserved regulatory signatures provides insight into core gene regulatory mechanisms underlying basal ganglia function. This comparative visualization capability allows users to assess both conserved and species-specific regulatory features within a single interactive framework, supporting evolutionary and translational analyses of basal ganglia gene regulation.

**Figure 2.**
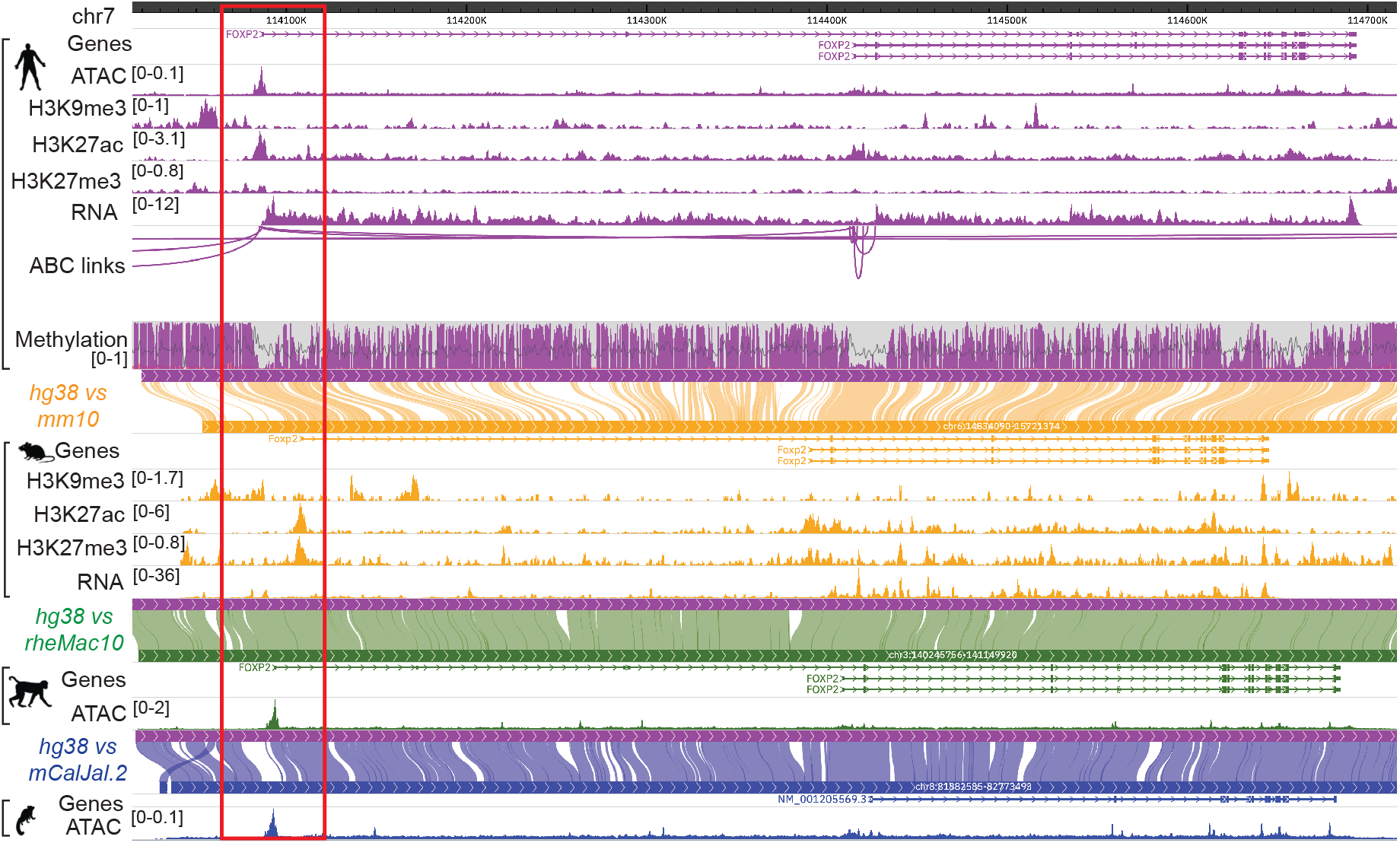
Cross-species comparative epigenomic visualization at the *FOXP2* locus of the striatal hybrid D1D2 hybrid medium spiny neurons (STR D1D2 Hybrid MSNs). Genome browser view showing epigenomic datasets from human, mouse, macaque, and marmoset aligned at the *FOXP2* gene locus. Tracks are color-coded by species: human (purple), mouse (orange), macaque (green), and marmoset (blue). Conserved open chromatin and putative enhancer signatures across species are highlighted, with red boxes indicating regions exhibiting shared regulatory activity in all four species. Cross-species alignment is enabled by genomeAlign tracks, generated from pairwise whole-genome alignments between each species, allowing homologous regulatory elements to be visualized in a unified coordinate framework. (xenoRefGene annotation for marmoset was downloaded from UCSC Genome hub^21^: GCA_011100555.2 genome assembly, all other species used refGene annotation from UCSC Genome Browser^22^)

### Single cell multi-Omics data analysis platform

In addition to genome browser-based visualization, the portal provides integrated tools for exploring single-cell and single-nucleus analysis results through commonly used summary and dimensionality reduction plots^23^. This module (scAnalysis) allows users to examine molecular variation across cells, regions, and cell types without requiring external computational workflows (Figure 3). Users can flexibly select datasets, assay modalities, and features of interest, including predefined metadata variables, individual genes, or gene lists for analysis.

**Figure 3.**
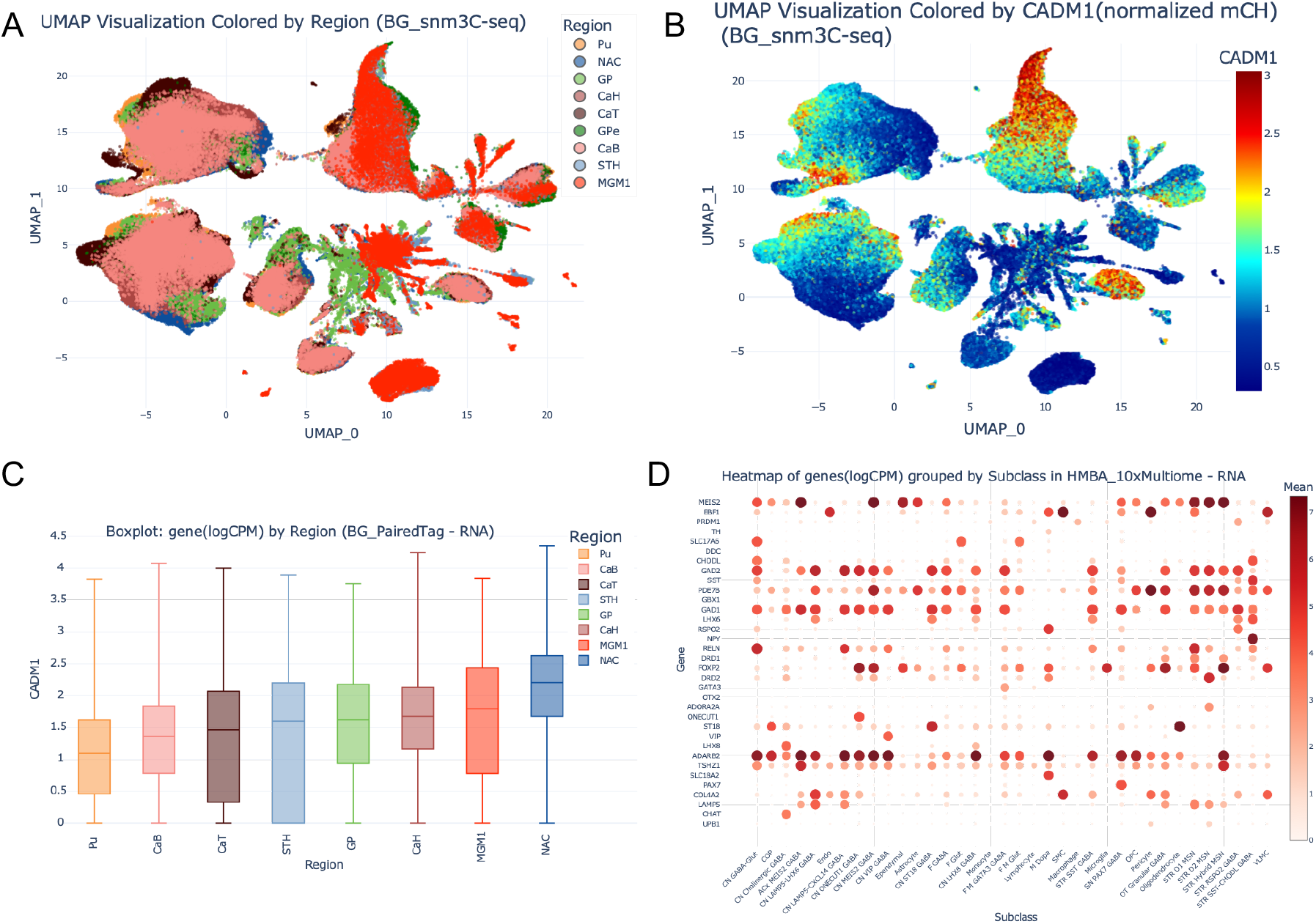
Visualization of single-cell analysis results within the portal. The portal supports interactive visualization of single-cell and single-nucleus analysis results using dimensionality reduction, distribution, and summary plots. Users first select a dataset and assay modality, then generate plots based on predefined variables, individual gene, or gene sets. (A) UMAP embedding of single nuclei from snm3C-seq data, colored by basal ganglia subregions, including head of caudate (CaH), body of caudate (CaB), tail of caudate (CaT), putamen (Pu), nucleus accumbens (NAC), globus pallidus (GP), subthalamic nucleus (STH) and gray matter of midbrain (MGM1). The UMAP interface also supports side-by-side plotting for comparative visualization. (B) The same UMAP embedding but colored by the CH methylation of gene CADM1. (C) Boxplot showing *CADM1* gene expression distributions across basal ganglia regions using RNA expression data from Paired-Tag. (D) Dot heatmap displaying average expression levels for a user-defined cell-type specific marker genes in basal ganglia, grouped by subclass. Rows correspond to genes and columns correspond to cell subclasses.

As an example, a UMAP visualization of snm3C-seq data can be used to examine cell-type-specific DNA methylation patterns at genes of interest. *CADM1* encodes a synaptic cell adhesion molecule that plays important roles in neuronal connectivity and synapse formation and has been implicated in neurodevelopmental and neuropsychiatric disorders, making it a biologically relevant locus for comparative analysis^24,25^. When coloring UMAP embeddings by normalized CH methylation (Ding et al., co-submit) levels at the *CADM1* locus, which was reported to show gradient pattern from anterior NAC, CaH to CaB and posterior CaT in gene expression, DNA methylation and 3D chromatin interaction. The same gradient can be easily reproduced in the scAnalysis module by querying region (Figure 3A) and gene *CADM1* (Figure 3B). The ability to display two UMAP plots side by side further facilitates direct comparison across genes, modalities, or experimental conditions (Supplementary Figure 1). The portal also supports region-level comparisons of gene expression using interactive boxplot. Visualization of *CADM1* expression across basal ganglia regions using Paired-Tag RNA data confirmed the high expression level of *CADM1* in NAC (Figure 3C). Such coordinated views enable users to relate epigenetic regulation to transcriptional outcomes within a unified interface. Finally, gene set-based analyses are supported through heatmap visualizations that summarize epigenomic signals across basal ganglia cell types. For a user-defined gene list, expression or DNA methylation from different basal ganglia datasets can be displayed and grouped by subclass, enabling identification of cell-type-specific signatures (Figure 3D). Together, these single-cell visualization tools extend the portal beyond genomic coordinate-centric views and provide a flexible framework for interrogating regulatory and transcriptional heterogeneity in the basal ganglia.

### Example biological insights enabled by integrated basal ganglia multi-omics analyses

To illustrate the utility of the resource for biological discovery, we present representative examples demonstrating how integrated multi-omics visualization across cell types can reveal regulatory features of the basal ganglia. These examples are intended to highlight analytical workflows enabled by the platform rather than to provide a comprehensive biological interpretation.

#### Cell-type-specific enhancer signature

To illustrate the portal’s ability to resolve cell-type-specific regulatory elements, we examined open chromatin accessibility at the *DRD2* locus using 10x Multiome ATAC-seq data (Figure 4). This analysis reveals an enhancer verified by AAV toolbox assay^26^ that is selectively accessible in cerebral nuclei (CN) Lateral Ganglionic Eminence (LGE)-derived GABAergic neurons^27^, including medium spiny neurons (MSN)^28^, but not in non-neuronal cell populations. The cell-type-restricted accessibility pattern suggests that this enhancer may contribute to the selective regulation of *DRD2* expression in basal ganglia projection neurons. Such examples demonstrate how integrated single-cell chromatin accessibility visualization enables the identification of cell-type-specific candidate regulatory elements, providing mechanistic insight into the transcriptional programs underlying basal ganglia circuit specialization.

**Figure 4.**
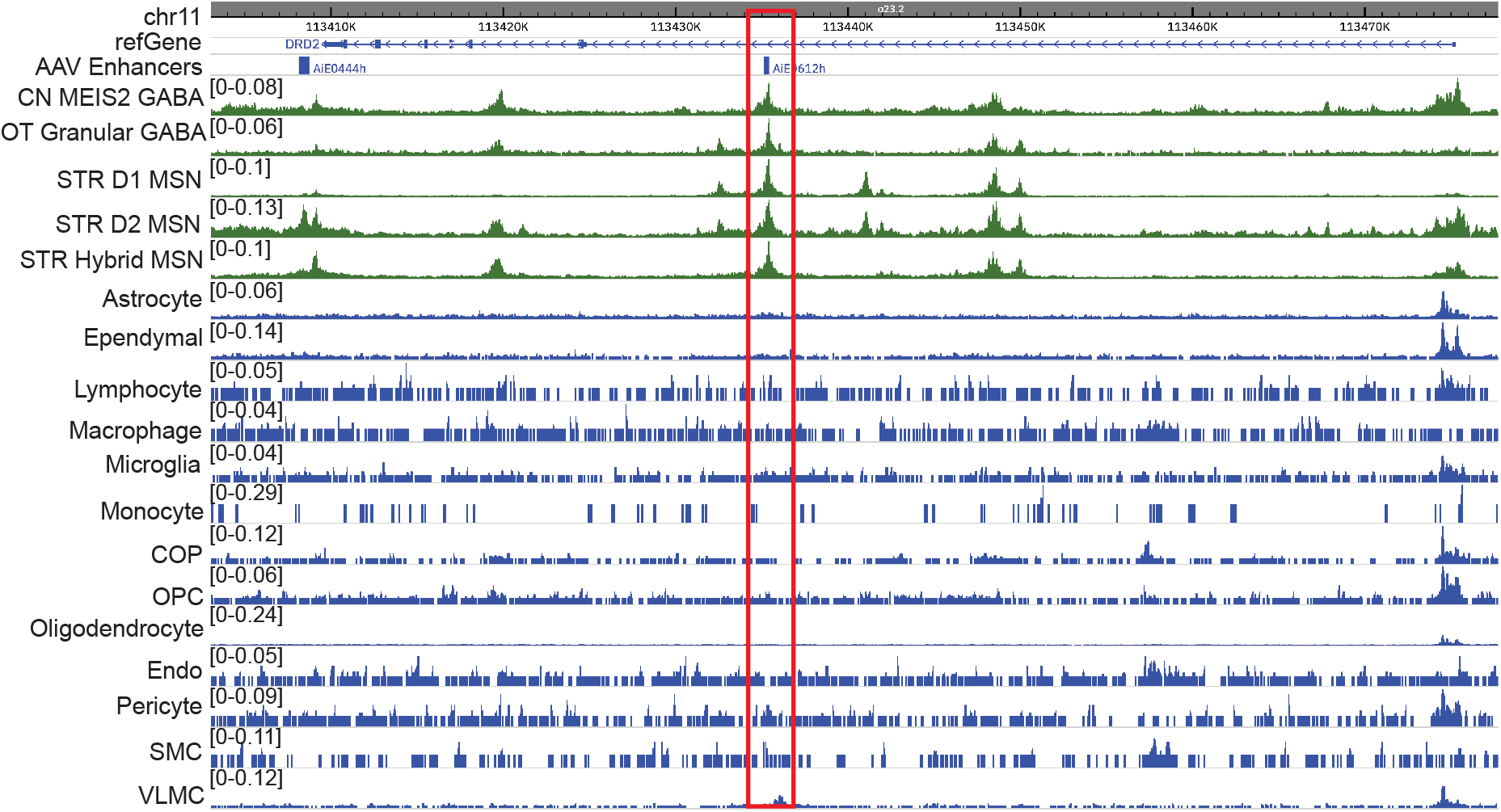
Cell-type-specific open chromatin signatures at a basal ganglia enhancer. Genome browser visualization of open chromatin profiles derived from 10x Multiome ATAC-seq data at the *DRD2* locus on chromosome 11. An enhancer exhibits strong accessibility selectively in CN LGE-derived GABAergic neurons, while remaining inactive in non-neuronal cell types. From top to bottom, tracks display the genomic coordinate ruler, gene annotations including *DRD2*, and annotated AAV enhancer elements shown as blocks beneath the gene model. All tracks represent ATAC-seq signal (CPM), with green tracks corresponding to CN LGE-derived GABAergic neurons and blue tracks representing non-neuronal cells. The red box highlights an enhancer region with enriched chromatin accessibility specifically in MSNs and two additional GABAergic neuron subtypes.

#### Integration of chromatin conformation with epigenomic and transcriptional data

To demonstrate how integrated multi-omics data can reveal cell-type-specific regulatory mechanisms, we examined chromatin interactions and transcriptional activity across neuronal and non-neuronal populations using combined snm3C-seq, ATAC-seq, and Paired-Tag datasets (Figure 5). The portal enables simultaneous visualization of chromatin accessibility, histone modifications, three-dimensional chromatin contacts, and gene expression, providing a comprehensive view of regulatory architecture within defined cell types. At the *BCL11B* locus, one of the MSN marker genes^29^, striatal MSNs exhibit increased Activity-By-Contact (ABC) model^30,31^ predicted enhancer-promoter interactions (Ding et al., co-submit) accompanied by elevated RNA expression relative to microglia. This illustrates how ABC modeling, when integrated with chromatin conformation and epigenomic data, can identify putative regulatory links underlying cell-type-specific transcriptional programs. By enabling direct contrast of regulatory interactions and expression across neuronal and non-neuronal cell types, the portal facilitates mechanistic investigation of gene regulation in the basal ganglia and supports the prioritization of candidate regulatory elements for functional follow-up studies. This also underscores the importance of higher-order chromatin structure in shaping basal ganglia gene expression programs and demonstrates the added value of incorporating chromatin conformation data into integrative analyses.

**Figure 5.**
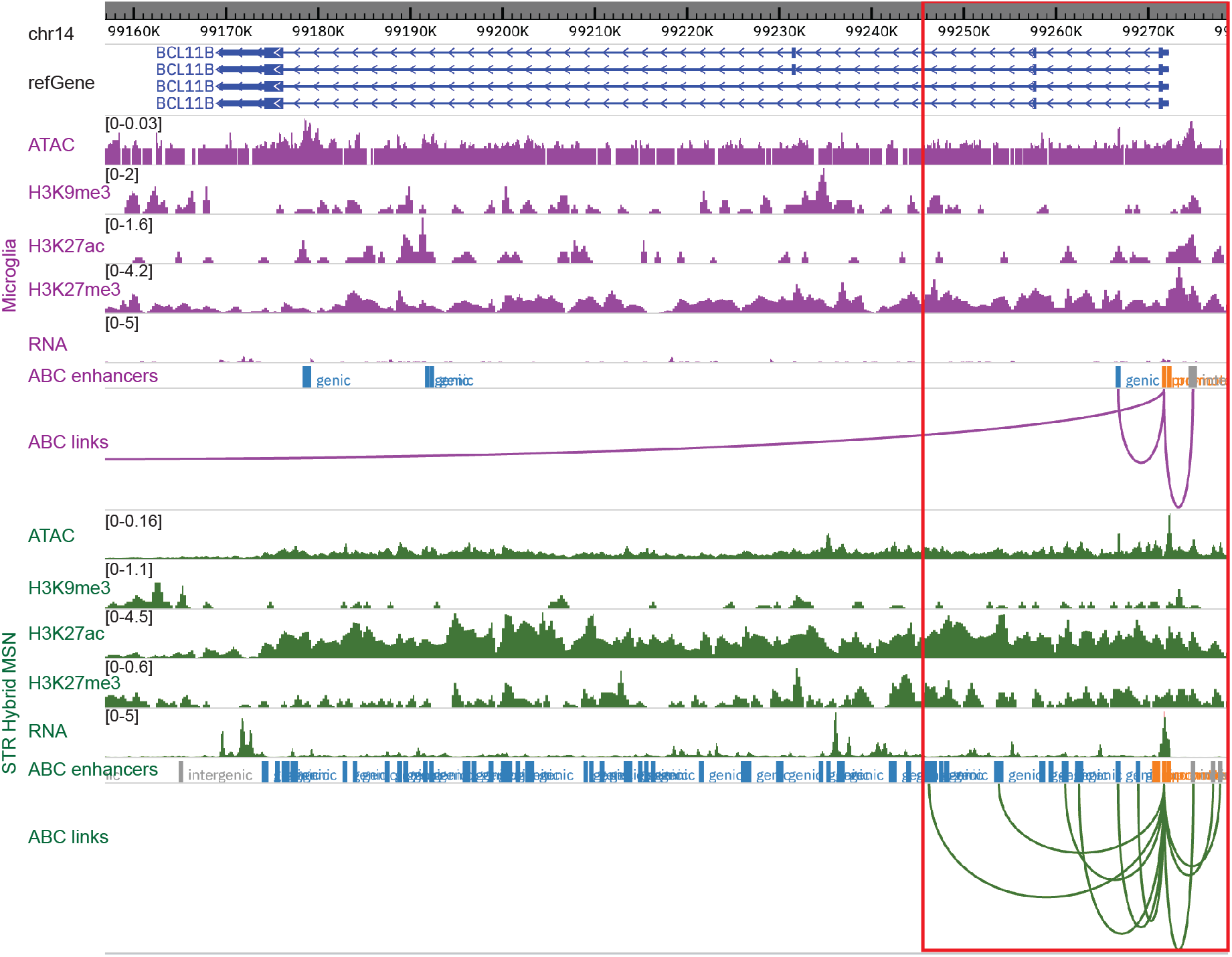
Cell-type-specific chromatin interactions associated with transcriptional differences in neuronal and non-neuronal populations. Genome browser view illustrates coordinated chromatin interaction and transcriptional profiles across the MSN marker gene *BCL11B* locus. Tracks include genomic coordinates, refGene annotations, 10x Multiome ATAC-seq signal, histone modification profiles and RNA expression measured by Paired-Tag, and activity-by-contact (ABC) regulatory links inferred from integrated snm3C-seq and ATAC-seq data. Purple tracks represent microglia, while green tracks represent striatal hybrid medium spiny neurons (STR Hybrid MSNs). The view highlights the *BCL11B* locus, where increased ABC interactions and higher RNA expression are observed in MSNs.

#### Cross-species conservation and divergence of regulatory elements

We observed that using the portal’s comparative epigenomics functionality, users can examine conserved regulatory features between different species (Figure 2). We can also examine the divergence, here using ventral striatum (STRv) D1 MSN as an example, aligned human and mouse datasets reveal distinct regulatory architectures at the *PLXND1* locus (Figure 6). While mouse D1 MSNs exhibit a pronounced enhancer signature characterized by H3K27ac enrichment accompanied by detectable gene expression, these features are substantially diminished or absent in the corresponding human cell population. Such analyses underscore the importance of human-specific epigenomic data for interpreting regulatory mechanisms in the basal ganglia and caution against direct extrapolation from model organisms without comparative validation.

**Figure 6.**
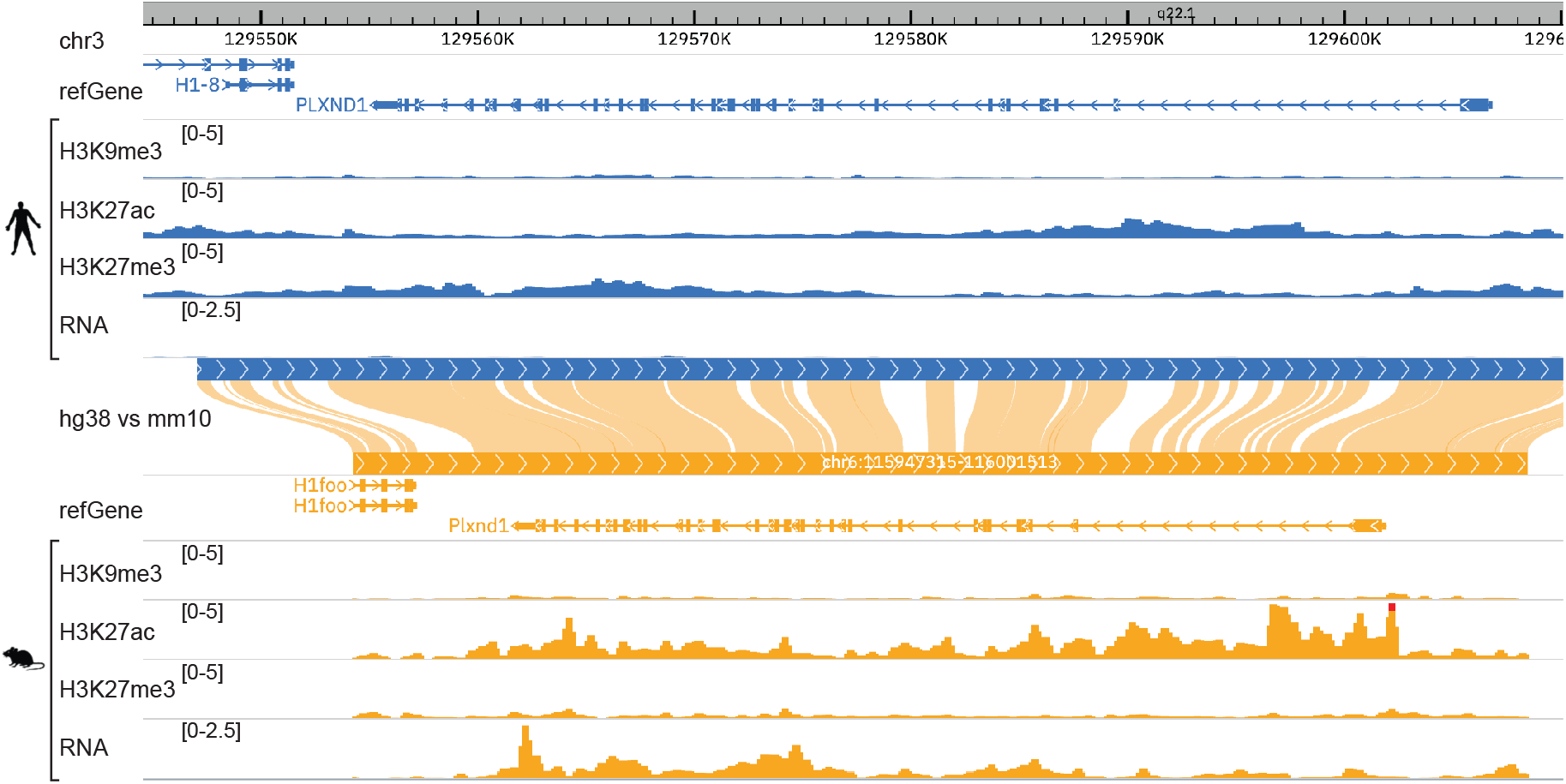
Species-specific epigenomic regulation at the PLXND1 locus in striatal D1 medium spiny neurons. Genome browser view showing aligned human and mouse epigenomic datasets from STRv D1 medium spiny neurons (MSNs) at the *PLXND1* gene locus. Comparative visualization reveals a mouse-specific enhancer signature marked by H3K27ac enrichment and corresponding RNA expression, whereas these regulatory and transcriptional signals are absent or markedly reduced in the human data. This example highlights species-specific differences in epigenomic regulation within homologous basal ganglia cell types.

## Discussion

In this study, we present an integrated, web-based resource for exploring the multi-omics regulatory landscape of the basal ganglia at both pseudo-bulk and single-cell resolution. By unifying transcriptomic, chromatin accessibility, DNA methylation, histone modification, and 3D chromatin conformation data across cell types, regions, and species, this platform addresses a critical need for accessible and interoperable tools to study the molecular architecture of basal ganglia circuits.

The example analyses presented here illustrate how coordinated visualization of multiple regulatory layers can provide mechanistic insights that are difficult to obtain from individual modalities alone. Cell-type-specific patterns of chromatin accessibility, DNA methylation, and histone modifications observed at key basal ganglia genes reinforce the concept that epigenetic regulation is a primary driver of neuronal identity and functional specialization. Integration of these epigenomic signals with gene expression data enables users to link regulatory features to transcriptional outcomes directly, facilitating hypothesis generation about gene regulation in defined neuronal populations.

Beyond genome browser-centric views, the platform’s single-cell analysis visualizations further enhance interpretation of regulatory heterogeneity. Dimensionality reduction, distribution, and gene set-based summary plots enable users to examine variation in transcriptional and epigenetic features across individual cells, cell types, brain regions, and other metadata. For example, UMAP visualizations colored by gene-specific methylation or expression values reveal cell-type-specific regulatory patterns that are not readily apparent from aggregated tracks alone, while boxplots and heatmaps summarize region- and subclass-level differences across genes or gene sets. These single-cell views complement coordinate-based visualization by providing a cell-centric perspective on regulatory diversity within the basal ganglia.

Importantly, incorporation of chromatin conformation data from snm3C-seq adds a structural dimension to these analyses, enabling examination of long-range regulatory interactions in a cell-type-specific manner. Combined with single-cell methylation and expression analyses, these data highlight how three-dimensional genome organization, epigenetic state, and transcriptional output jointly shape basal ganglia gene regulation. This integrated framework is particularly relevant for interpreting non-coding regulatory elements and disease-associated variants that act by disrupting regulatory architecture rather than through protein-coding sequences.

Cross-species comparative analyses further underscore the value of this resource. While many regulatory features are conserved between human and mouse basal ganglia cell types, species-specific differences in non-coding regulatory elements are readily apparent. The ability to examine these features both at genomic loci and through single-cell summaries reinforces the importance of human-specific epigenomic data for interpreting genetic risk loci and cautions against over-generalization from model organisms alone. Coordinated visualization of homologous regions across species enables systematic investigation of conserved and divergent regulatory mechanisms.

Beyond biological insights, this work emphasizes the importance of data accessibility and usability. The web portal’s modular architecture, interactive visualization framework, and integration with the WashU Epigenome Browser lower technical barriers for researchers with diverse computational backgrounds. Features such as multi-species track views, interactive single-cell plots, and shareable visualization states support reproducible, collaborative exploration and align with the open-science principles of the NIH BRAIN Initiative and BICAN.

As single-cell and multimodal profiling technologies continue to evolve, this resource provides a flexible foundation for incorporating additional datasets, brain regions, and species. Future expansions may include additional data from NHP species and extended single-cell analytical capabilities. Collectively, this platform serves as a community-oriented foundation for advancing our understanding of basal ganglia biology and its role in neurological and psychiatric disease.

## Conclusion

In summary, we present an open, interactive platform that integrates single-cell transcriptomic and epigenomic data to enable systematic exploration of the basal ganglia at cellular and regulatory resolution. In addition to genome-centric views, the platform provides dedicated single-cell analysis visualizations that capture regulatory and transcriptional heterogeneity across cell types, regions, and gene sets. By combining complementary modalities with intuitive visualization and comparative tools, this resource lowers barriers to data access and supports integrative investigations of gene regulation, genome organization, and cell-type specialization. As part of BICAN infrastructure studies, this platform serves as a foundational reference for basal ganglia epigenome research and a scalable framework for future expansion to advance our understanding of brain function and disease.

## Acknowledgements

We are deeply grateful to the scientists at the Allen Institute who developed the consensus basal ganglia cell taxonomy, and the anonymous donors who contributed the brain samples.

## Supplementary Figures

**Supplementary Figure 1.**
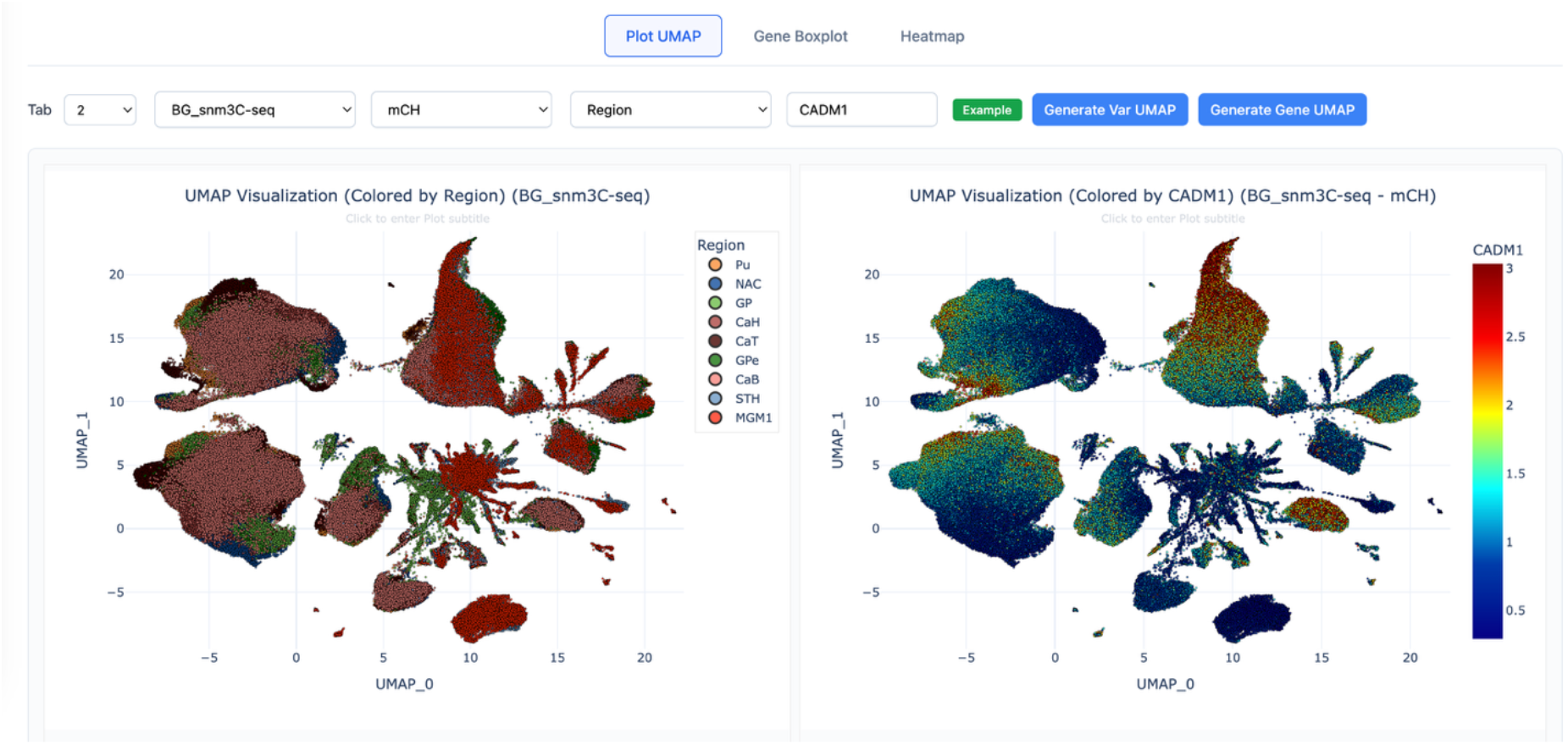
Multiple UMAP plots in the single cell analysis visualization module allows easier side-by-side comparison.

