## Supplementary_information for "An Integrated Single-Cell and Epigenomic Resource for Comparative Analysis of the Basal Ganglia"

Supplementary Table 1: Datasets organized by the subclass level of cell taxonomy.

| Neighborhood | Class | Subclass | 10X multiome (Human) | 10X multiome (Macaque) | 10X multiome (Marmoset) | Paired Tag (Human) | Paired Tag (Mouse) | snm3C-seq (Human) |
| --- | --- | --- | --- | --- | --- | --- | --- | --- |
| Glut Sero Dopa | F M Glut | F Glut | Y | N | N | Y | N | Y |
| Glut Sero Dopa | F M Glut | F M Glut | Y | N | N | Y | N | Y |
| Glut Sero Dopa | M Dopa | M Dopa | Y | N | N | Y | N | Y |
| Nonneuron | Astro-Epen | Astrocyte | Y | N | N | Y | N | Y |
| Nonneuron | Astro-Epen | Ependymal | Y | N | N | Y | N | N |
| Nonneuron | Immune | Lymphocyte | Y | N | N | Y | N | Y |
| Nonneuron | Immune | Macrophage | Y | N | N | Y | N | N |
| Nonneuron | Immune | Microglia | Y | N | N | Y | N | Y |
| Nonneuron | Immune | Monocyte | Y | N | N | Y | N | N |
| Nonneuron | OPC-Oligo | COP | Y | N | N | Y | N | N |
| Nonneuron | OPC-Oligo | OPC | Y | N | N | Y | N | Y |
| Nonneuron | OPC-Oligo | Oligodendrocyte | Y | N | N | Y | N | Y |
| Nonneuron | Vascular | Endo | Y | N | N | Y | N | Y |
| Nonneuron | Vascular | Pericyte | Y | N | N | Y | N | Y |
| Nonneuron | Vascular | SMC | Y | N | N | Y | N | Y |
| Nonneuron | Vascular | VLMC | Y | N | N | Y | N | Y |
| Subpallium GABA | CN CGE GABA | CN LAMP5-CXCL14 GABA | Y | N | N | Y | N | Y |
| Subpallium GABA | CN CGE GABA | CN VIP GABA | Y | N | N | Y | N | Y |
| Subpallium GABA | CN LGE GABA | CN MEIS2 GABA | Y | N | N | Y | N | Y |
| Subpallium GABA | CN LGE GABA | OT Granular GABA | Y | N | N | Y | N | Y |
| Subpallium GABA | CN LGE GABA | STR D1 MSN | Y | N | N | Y | N | Y |
| Subpallium GABA | CN LGE GABA | STR D2 MSN | Y | N | N | Y | N | Y |
| Subpallium GABA | CN LGE GABA | STR Hybrid MSN | Y | N | N | Y | N | Y |
| Subpallium GABA | CN MGE GABA | CN Cholinergic GABA | Y | N | N | Y | N | Y |
| Subpallium GABA | CN MGE GABA | CN LAMP5-LHX6 GABA | Y | N | N | Y | N | Y |
| Subpallium GABA | CN MGE GABA | CN ST18 GABA | Y | N | N | Y | N | Y |
| Subpallium GABA | CN MGE GABA | STR RSPO2 GABA | Y | N | N | Y | N | Y |
| Subpallium GABA | CN MGE GABA | STR SST GABA | Y | N | N | Y | N | N |
| Subpallium GABA | CN MGE GABA | STR SST-CHODL GABA | Y | N | N | Y | N | Y |
| Subpallium GABA | Cx GABA | ACx MEIS2 GABA | Y | N | N | Y | N | Y |
| Subpallium GABA | F M GABA | CN LHX8 GABA | Y | N | N | Y | N | Y |
| Subpallium GABA | F M GABA | CN ONECUT1 GABA | Y | N | N | Y | N | Y |
| Subpallium GABA | F M GABA | F GABA | Y | N | N | Y | N | Y |
| Subpallium GABA | F M GABA | F M GATA3 GABA | Y | N | N | Y | N | Y |
| Subpallium GABA | F M GABA | SN PAX7 GABA | Y | N | N | Y | N | Y |
| Subpallium GABA-Glut | CN GABA-Glut | CN GABA-Glut | Y | N | N | Y | N | Y |

Supplementary Table 2: Datasets organized by the group level of cell taxonomy.

| Neighborhood | Class | Subclass | Group | 10X multiome (Human) | 10X multiome (Macaque) | 10X multiome (Marmoset) | Paired Tag (Human) | Paired Tag (Mouse) | snm3C-seq (Human) |
| --- | --- | --- | --- | --- | --- | --- | --- | --- | --- |
| Glut Sero Dopa | F M Glut | F Glut | BF SKOR1 Glut | Y | Y | Y | Y | N | Y |
| Glut Sero Dopa | F M Glut | F Glut | STH PVALB-PITX2 Glut | Y | Y | Y | Y | N | Y |
| Glut Sero Dopa | F M Glut | F M Glut | VTR-HTH Glut | Y | Y | Y | Y | N | Y |
| Glut Sero Dopa | M Dopa | M Dopa | SN SOX6 Dopa | Y | Y | Y | Y | N | Y |
| Glut Sero Dopa | M Dopa | M Dopa | SN-VTR CALB1 Dopa | Y | Y | Y | Y | N | Y |
| Glut Sero Dopa | M Dopa | M Dopa | SN-VTR GAD2 Dopa | Y | Y | Y | Y | N | Y |
| Nonneuron | Astro-Epen | Astrocyte | Astrocyte | Y | Y | Y | Y | N | N |
| Nonneuron | Astro-Epen | Astrocyte | Astrocyte-1 | N | N | N | N | N | Y |
| Nonneuron | Astro-Epen | Astrocyte | Astrocyte-2 | N | N | N | N | N | Y |
| Nonneuron | Astro-Epen | Astrocyte | Astrocyte-3 | N | N | N | N | N | Y |
| Nonneuron | Astro-Epen | Astrocyte | Astrocyte-4 | N | N | N | N | N | Y |
| Nonneuron | Astro-Epen | Astrocyte | ImAstro | Y | Y | N | Y | N | N |
| Nonneuron | Astro-Epen | Ependymal | Ependymal | Y | Y | Y | Y | N | N |
| Nonneuron | Immune | Lymphocyte | B cells | Y | N | N | N | N | N |
| Nonneuron | Immune | Lymphocyte | Lymphocyte | N | N | N | N | N | Y |
| Nonneuron | Immune | Lymphocyte | T cells | Y | Y | N | Y | N | N |
| Nonneuron | Immune | Macrophage | BAM | Y | Y | Y | Y | N | N |
| Nonneuron | Immune | Microglia | Microglia | Y | Y | Y | Y | N | N |
| Nonneuron | Immune | Microglia | Microglia-1 | N | N | N | N | N | Y |
| Nonneuron | Immune | Microglia | Microglia-2 | N | N | N | N | N | Y |
| Nonneuron | Immune | Microglia | Microglia-3 | N | N | N | N | N | Y |
| Nonneuron | Immune | Monocyte | Monocyte | Y | Y | N | Y | N | N |
| Nonneuron | OPC-Oligo | COP | COP | Y | Y | Y | Y | N | N |
| Nonneuron | OPC-Oligo | OPC | OPC | Y | Y | Y | Y | N | Y |
| Nonneuron | OPC-Oligo | Oligodendrocyte | ImOligo | Y | Y | N | Y | N | N |
| Nonneuron | OPC-Oligo | Oligodendrocyte | Oligo OPALIN | Y | Y | N | Y | N | Y |
| Nonneuron | OPC-Oligo | Oligodendrocyte | Oligo PLEKHG1 | Y | Y | Y | Y | N | Y |
| Nonneuron | Vascular | Endo | Endo | Y | Y | Y | Y | N | Y |
| Nonneuron | Vascular | Pericyte | Pericyte | Y | Y | Y | Y | N | Y |
| Nonneuron | Vascular | SMC | SMC | Y | Y | Y | Y | N | Y |
| Nonneuron | Vascular | VLMC | VLMC | Y | Y | Y | Y | N | Y |
| Subpallium GABA | CN CGE GABA | CN LAMP5-CXCL14 GABA | LAMP5-CXCL14 GABA | Y | Y | Y | Y | N | Y |
| Subpallium GABA | CN CGE GABA | CN VIP GABA | VIP GABA | Y | Y | Y | Y | N | Y |
| Subpallium GABA | CN LGE GABA | CN MEIS2 GABA | GPe MEIS2-SOX6 GABA | Y | Y | Y | Y | N | Y |
| Subpallium GABA | CN LGE GABA | OT Granular GABA | OT D1 ICj | Y | Y | Y | Y | N | Y |
| Subpallium GABA | CN LGE GABA | STR D1 MSN | STRd D1 Matrix MSN | Y | Y | N | Y | Y | Y |
| Subpallium GABA | CN LGE GABA | STR D1 MSN | STRd D1 Striosome MSN | Y | Y | Y | Y | Y | Y |
| Subpallium GABA | CN LGE GABA | STR D1 MSN | STRv D1 MSN | Y | Y | Y | Y | Y | Y |
| Subpallium GABA | CN LGE GABA | STR D2 MSN | STRd D2 Matrix MSN | Y | Y | Y | Y | Y | Y |
| Subpallium GABA | CN LGE GABA | STR D2 MSN | STRd D2 StrioMat Hybrid MSN | Y | Y | Y | Y | Y | Y |
| Subpallium GABA | CN LGE GABA | STR D2 MSN | STRd D2 Striosome MSN | Y | Y | N | Y | Y | Y |
| Subpallium GABA | CN LGE GABA | STR D2 MSN | STRv D2 MSN | Y | Y | Y | Y | Y | Y |
| Subpallium GABA | CN LGE GABA | STR Hybrid MSN | STR D1D2 Hybrid MSN | Y | Y | Y | Y | Y | Y |
| Subpallium GABA | CN LGE GABA | STR Hybrid MSN | STRv D1 NUDAP MSN | Y | Y | Y | Y | Y | Y |
| Subpallium GABA | CN MGE GABA | CN Cholinergic GABA | GPin-BF Cholinergic GABA | Y | Y | Y | Y | N | Y |
| Subpallium GABA | CN MGE GABA | CN Cholinergic GABA | STR Cholinergic GABA | Y | Y | Y | Y | N | Y |
| Subpallium GABA | CN MGE GABA | CN Cholinergic GABA | STRd Cholinergic GABA | Y | Y | Y | Y | N | Y |
| Subpallium GABA | CN MGE GABA | CN LAMP5-LHX6 GABA | LAMP5-LHX6 GABA | Y | Y | Y | Y | N | Y |
| Subpallium GABA | CN MGE GABA | CN ST18 GABA | STR FS PTHLH-PVALB GABA | Y | Y | N | Y | N | Y |
| Subpallium GABA | CN MGE GABA | CN ST18 GABA | STR TAC3-PLPP4 GABA | Y | Y | Y | Y | N | Y |
| Subpallium GABA | CN MGE GABA | CN ST18 GABA | STR-BF TAC3-PLPP4-LHX8 GABA | Y | Y | Y | Y | N | Y |
| Subpallium GABA | CN MGE GABA | STR RSPO2 GABA | STR LYPD6-RSPO2 GABA | Y | Y | Y | Y | N | Y |
| Subpallium GABA | CN MGE GABA | STR RSPO2 GABA | STR SST-RSPO2 GABA | Y | Y | Y | Y | N | Y |
| Subpallium GABA | CN MGE GABA | STR SST GABA | STR SST-ADARB2 GABA | Y | Y | Y | Y | N | N |
| Subpallium GABA | CN MGE GABA | STR SST-CHODL GABA | STR SST-CHODL GABA | Y | Y | Y | Y | N | N |
| Subpallium GABA | CN MGE GABA | STR SST-CHODL GABA | STR SST-CHODL GRIN2A GABA | N | N | N | N | N | Y |
| Subpallium GABA | CN MGE GABA | STR SST-CHODL GABA | STR SST-CHODL NRP2 GABA | N | N | N | N | N | Y |
| Subpallium GABA | CN MGE GABA | STR SST-CHODL GABA | STR SST-CHODL RORA GABA | N | N | N | N | N | Y |
| Subpallium GABA | Cx GABA | ACx MEIS2 GABA | ACx MEIS2 GABA | N | N | N | N | N | Y |
| Subpallium GABA | Cx GABA | ACx MEIS2 GABA | OB Dopa-GABA | Y | Y | Y | Y | N | N |
| Subpallium GABA | Cx GABA | ACx MEIS2 GABA | OB FRMD7 GABA | Y | Y | Y | Y | N | N |
| Subpallium GABA | F M GABA | CN LHX8 GABA | GPe SOX6-CTXND1 GABA | Y | Y | Y | Y | N | Y |
| Subpallium GABA | F M GABA | CN LHX8 GABA | GPe-NDB-SI LHX6-LHX8-GBX1 GABA | Y | Y | Y | Y | N | Y |
| Subpallium GABA | F M GABA | CN ONECUT1 GABA | GPi Core | Y | Y | Y | Y | N | Y |
| Subpallium GABA | F M GABA | F GABA | AMY-SLEA-BNST D1 GABA | Y | Y | N | Y | N | N |
| Subpallium GABA | F M GABA | F GABA | AMY-SLEA-BNST GABA | Y | Y | Y | Y | N | Y |
| Subpallium GABA | F M GABA | F GABA | F GABA | N | N | N | N | N | Y |
| Subpallium GABA | F M GABA | F GABA | ZI-HTH GABA | Y | Y | Y | Y | N | Y |
| Subpallium GABA | F M GABA | F M GATA3 GABA | F M GATA3 GABA | N | N | N | Y | N | Y |
| Subpallium GABA | F M GABA | F M GATA3 GABA | SN GATA3-PVALB GABA | Y | Y | Y | Y | N | Y |
| Subpallium GABA | F M GABA | F M GATA3 GABA | SN-VTR-HTH GATA3-TCF7L2 GABA | Y | Y | Y | Y | N | Y |
| Subpallium GABA | F M GABA | SN PAX7 GABA | SN EBF2 GABA | Y | N | Y | Y | N | Y |
| Subpallium GABA | F M GABA | SN PAX7 GABA | SN SEMA5A GABA | Y | N | Y | Y | N | Y |
| Subpallium GABA-Glut | CN GABA-Glut | CN GABA-Glut | GPi Shell | Y | Y | Y | Y | N | Y |

Supplementary file 1: Complete list of available data in the TSV (Tab-Separated Values) format.
